# Fusion dynamics and size-dependent droplet microstructure in ssDNA mediated protein phase separation

**DOI:** 10.1101/2023.11.13.566798

**Authors:** Yunqiang Bian, Wenfei Li

## Abstract

Biomolecular cocondensation involving proteins and nucleic acids has been recognized to play crucial roles in genome organization and transcriptional regulation. However, the biophysical mechanisms underlying the fusion dynamics and microstructure evolution of the droplets during the early stage of liquid-liquid phase separation (LLPS) remain elusive. In this work, we study the phase separation of linker histone H1, which is among the most abundant chromatin proteins, in the presence of single-stranded DNA (ssDNA) capable of forming G-quadruplex structures by using residue-resolved molecular dynamics simulations. Firstly, we uncovered a kinetic bottleneck step in the droplet fusion. Productive fusion events are triggered by the formation of ssDNA mediated electrostatic bridge within the contacting zone of two droplets. Secondly, the simulations revealed a size-dependence of the droplet microstructure and stoichiometry. With droplet growth, its microstructure evolves as driven by the maximization of the electrostatic contacts between ssDNA and the highly charged segment of H1. Finally, we showed that the folding of ssDNA to G-quadruplex promotes LLPS by increasing the multivalency and strength of protein-DNA interactions. These findings provided new mechanistic insights into the microstructure and growth dynamics of the biomolecular droplets formed during the early stage of the ssDNA-protein cocondensation.

## Introduction

Biological condensates formed by liquid-liquid phase separation (LLPS), in which biomolecules in solutions spontaneously separate into coexisting dilute and dense phases, are considered to play essential roles in myriad cellular processes, such as the stress responses, ribosome biogenesis, DNA damage repair, and chromatin organization.^1–19^ These membrane-less condensates not only provide a favorable environment for biomolecules to establish interaction networks efficiently, but also contribute to homeostasis through concentration buffering effect. Aberrant condensation, on the other hand, can be involved in the pathogenesis of numerous neurodegenerative diseases. Therefore, unraveling the key physiochemical factors that regulate the structure and dynamics of biological condensates is crucial for understanding the biophysical basis underlying the biologically and pathologically relevant molecular events.^20–27^

It has been well established that multivalent weak interactions play a central role in driving LLPS.^4^ Proteins containing intrinsically disordered regions (IDRs) of low-sequence complexity possess inherent multivalency and are widely engaged in the formation of membraneless condensates.^28,29^ Such weak and multivalent interactions renders the biological condensates dynamic and exchangeable with surrounding cellular environment. Thus the structure and dynamics of the condensates can be readily altered by various physiochemical factors or stimuli, such as pH, salt concentration, molecular crowding, protein folding/ligand binding, and post-translational modifications, which can potentially contribute to the regulation of cellular processes.^30–46^ The phase behavior can be further enriched by sequence segregation of IDRs, resembling the block copolymer, and by heterotypic inter-molecule interactions of multicomponent systems, which can often lead to the microphase separation and layered structures of condensates.^47^

Nucleic acid molecules, which are highly charged, are key inducers or regulators of biological condensation mainly through inter-molecule Coulombic interactions.^11,12,18,48–54^ Because of the complex phase behavior and their biological prevalence, such multi-component systems containing DNA and RNA received increasing attentions. A well documented example is that the presence of RNA molecules can promote the LLPS of fused in sarcoma (FUS), which has a RNA-binding domain and a low-complexity domain.^15,53,54^ In addition, both experimental and computational studies showed that cocondenation of protein and DNA/RNA tends to form layered multiphasic microstructure. Recently, the significance of single-stranded DNA (ssDNA) as a crucial mediator in diverse phase separation processes of proteins has been increasingly appreciated.^32,48,55–61^ ssDNA has been involved in nearly all genomic processes, such as transcription, replication, and DNA damage repair. Through specific or nonspecific recognition and binding interactions with proteins, they can promote phase separation and facilitate the assembly of protein condensates. In a recent experimental report, Leicher and coworkers found that ssDNA can mediate enhanced condensation of linker histone H1, which is among the most abundant chromatin binding proteins.^58^ The ssDNA induced H1 condensation can also be observed *in vivo* around replication forks, which is likely involved in the DNA damage responses. Although these pioneer studies on the ssDNA-regulated protein phase separation have provided significant new insights into the molecular mechanism of DNA-protein co-condensation, several important questions regarding the growth dynamics and microstructure of the ssDNA-protein condensates remain unclear, including: i) what physical interactions dictate the fusion of liquid droplet; ii) how the droplet microstructure and stoichiometry evolve with fusion at the early stage of condensation; and iii) how the ssDNA folding regulates the ssDNA-protein cocondensation.

In this study, we address the above questions by investigating the guanine-rich ssDNA induced condensation of the linker histone H1 using residue-resolved molecular dynamics simulations. Guanine-rich DNA sequences have a tendency to adopt unique structural arrangements called G-quadruplexes (G4s), which have been found to play crucial roles in diverse biological processes, such as regulation of gene expression and maintenance of genome integrity.^62^ It was considered as potential target of drug design for cancer therapy.^63^ Since G4-forming sequences are highly prevalent in heterochromatin regions, the cocondensation between DNA quadruplex and the linker histone H1 can be involved in the chromatin condensation and gene expression regulation.^64^ In addition, it also provides a model system for investigating the general mechanism underlying the molecular process of ssDNA induced phase separation. In a recent experimental report, Mimura *et al* showed that G4-ssDNA folding tends to promote the growth of the ssDNA-H1 condensate and slow down the molecular motility within the droplet.^61^ The importance of G4 structure in inducing the droplet formation in cell was also demonstrated by using biomimetic protocells, which showed a temperature-dependent reversible microphase separation of guanine-rich ssDNA and its binding protein.^65^ Despite its biological significance, computational studies of the G4 coupled phase separation is still lacking to our best knowledge. By performing molecular dynamics simulations, we can directly identify the key physical interactions contributing to the droplet fusion and the formation of microstructure in the condensates, which is otherwise challenging in experimental studies. We showed that the surface of droplet formed by the ssDNA and H1 is occupied by the modestly charged N-terminal domain (NTD) and globular domain (core domain), whereas the highly charged C-terminal domain (CTD) are deeply buried inside the droplet together with ssDNA. The fusion of droplets is triggered by the formation of ssDNA-bridge, in which one ssDNA molecule simultaneously forms electrostatic contacts with the positively charged H1 core domains from the surfaces of two fusing droplets, demonstrating the crucial role of kinetic control in droplet fusion. We also revealed a size dependence of the microstructure of the droplets, which is dictated by the maximization of the charge-charge interactions between the CTD and ssDNA. In addition, we showed that the folding of the ssDNA to G4 structure increases the local charge density and contributes to the enhanced LLPS. These results provide new insights into the general biophysical mechanism underlying the ssDNA coupled biological phase separation.

## Results

### Growth of condensate by coalescence of smaller clusters

Linker histone H1 is a typlical IDR protein. It consists of a short unstructured NTD, a conserved core domain and a long disordered CTD (Figure 1A,B).^59^ The CTD is rich in lysine residues and therefore highly positively charged. The NTD and core domain are also positively charged but are much more moderate. For the ssDNA, we used the 22-nt oncogene c-myc promoter ssDNA sequence Pu22 (d[TGAGGGTGGGTAGGGTGGGTAAA]),^66^ which is known to fold to a parallel G4 topology (Figure 1A,C) and was observed to promote the phase separation of H1.^61^

**Figure 1.**
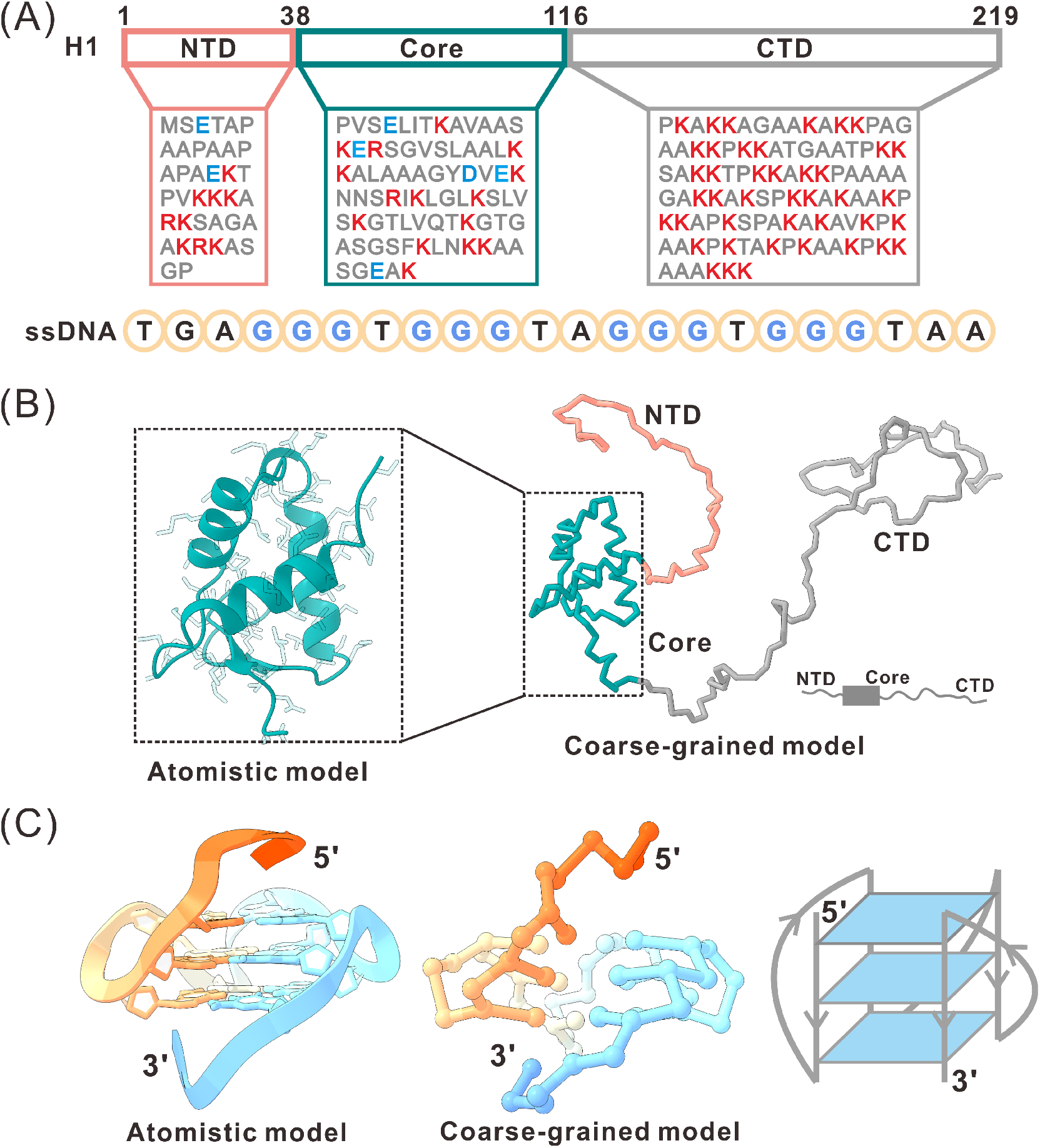
Simulation system. (A) Sequences of H1 and ssDNA (Pu22). Positively charged residues are colored red and negatively charged residues are colored blue. Guanine nucleotides forming the G4 structure are highlighted in blue. (B,C) The three-dimensional structures of H1 and G4 ssDNA used in the simulations. For each molecule, the atomistic model and the coarse-grained model are displayed, along with a schematic representation of the structure.

To obtain a comprehensive understanding of the phase separation dynamics, we performed 20 independent NVT simulations for a system comprising 60 H1 chains and 300 G4-ssDNA chains, a stoichiometry with the most prominent phase separation behavior as revealed by recent experimental study.^61^ The initial state of the system was prepared by randomly placing the biomolecules in a periodic cubic box with the size of 650*Å* × 650*Å* × 650 *Å*. Figure 2A shows a representative trajectory that effectively captured the transition of H1 chains from free monomers to larger condensates together with ssDNA molecules over time, which can provide an insightful understanding to the formation and growth dynamics of the droplet. Initially, the protein molecules exhibit a transition from individual monomers to various small clusters, indicating their tendency to immediately condense upon initialization. As the simulation progresses, these small clusters gradually assemble and coalesce, leading to the formation of larger condensates. Once formed, these larger condensates hardly dissolve, suggesting that Brownian coalescence,^67^ not Ostwald ripening,^68^ dominates the fusion events in the simulations.

**Figure 2.**
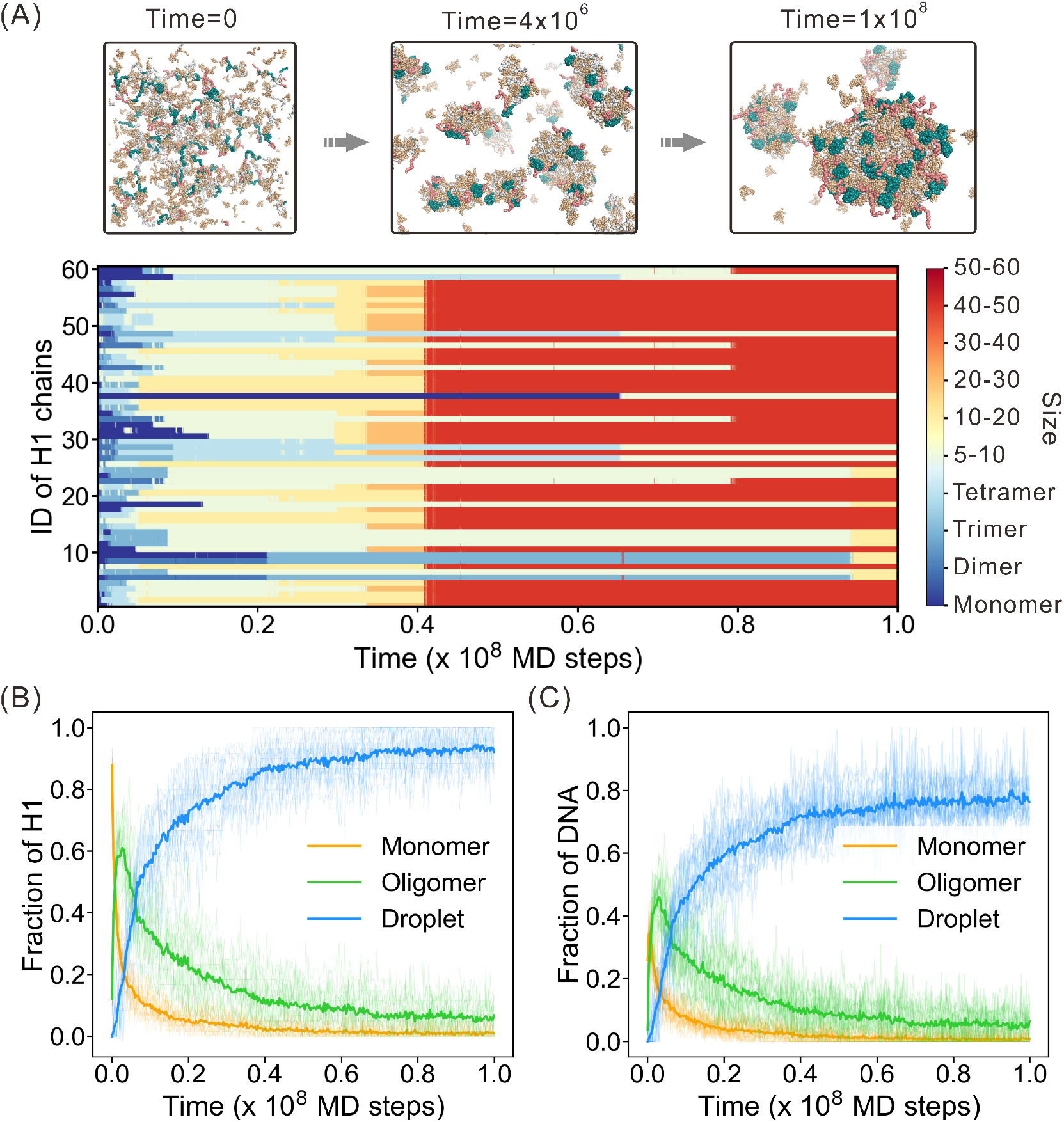
Molecular processes of the early stage of ssDNA-H1 cocondensation. (A) A representative simulation trajectory capturing the condensation of H1 under the regulation of G4-ssDNA. Representative snapshots of the MD simulation trajectory at different time points are shown in the upper panel. The cluster size to which each H1 chain belongs during the simulation process is shown in the lower panel. (B) Fraction of H1 in free monomers, oligomers and droplets as a function of simulation time. (C) Fraction of G4-ssDNA in free monomers, oligomers and droplets as a function of simulation time.

In further analysis for the overall 20 simulation trajectories, we categorized the assembled clusters into three distinct groups based on the number of constituent H1 chains (N*_H_*_1_), including monomers (N*_H_*_1_=1), oligomers (1*<*N*_H_*_1_*≤*4), and condensates (N*_H_*_1_*>*4). Throughout this work, we used N*_H_*_1_ to describe the droplet size. By adopting this classification, we were able to obtain a more comprehensive characterization of the condensation dynamics. Figure 2B illustrates the fraction of H1 chains presented in the above three classes of assembly clusters throughout the simulations. The fraction of H1 chains in the monomeric state was demonstrated to decrease monotonically over time. This is a direct consequence of the phase separation, as the H1 chains undergo condensation to form larger assemblies. Conversely, the fraction of H1 chains in droplet state was illustrated to increase monotonically. In contrast to the monotonic evolution of the monomeric state and droplet state, the fraction of H1 chains in oligomers exhibits a different pattern during the simulations. It reaches a peak value rapidly at the beginning of the simulations and then decreases as time progresses. Remarkably, the decrease in the fraction of H1 chains in oligomers coincides with the increase in the fraction of H1 chains in droplets. This observation supports the notion that the formation of larger droplets arises from the collision and fusion of smaller ones as suggested by the Brownian coalescence mechanism of droplet growth.^69^ The merging of smaller droplets contributes to the growth of larger and more stable structures. The time evolution of the fraction of ssDNA molecules in the three types of assemblies, as illustrated in Figure 2C, exhibited a similar pattern, which is a result of the ssDNA-H1 cocondensation.

### Key interactions contributing to the droplet fusion dynamics

The above simulations clearly demonstrated the importance of fusion of small clusters in the droplet growth. To gain a deeper understanding of the fusion dynamics involved in droplet growth, we conducted an additional set of ten independent simulations specifically focusing on the fusion process of two droplets (MD1). In constructing the system for fusion simulations, we selected a droplet (N*_H_*_1_=40) formed in the above simulations and then placed two copies with significant spatial separation in a cubic box (700*Å* x 700*Å* x 700 *Å*). In the above selected droplet, the molar ratio between ssDNA and H1 is lower than the average molar ratio (5.0) of the initial simulation system. To maintain the overall molar ratio intact, additional free G4-ssDNA molecules were randomly added in the fusion simulations. Figure 3A shows the instantaneous conformations at various time points in a representative trajectory that successfully captured the fusion event. For this trajectory, we also monitored the shape change of the system formed by the two component droplets over time by estimating the aspect ratio. A significant decrease in aspect ratio was observed in the fusion simulations, indicating a transformation from separated droplets to a more spherical, fused droplet. This observation demonstrates that the simulations can well capture the main feature of the fusion events. Overall, out of the ten trajectories examined, six successfully captured the full fusion event between two droplets within a simulation length of 5×10^7^ MD steps (Figure S1).

**Figure 3.**
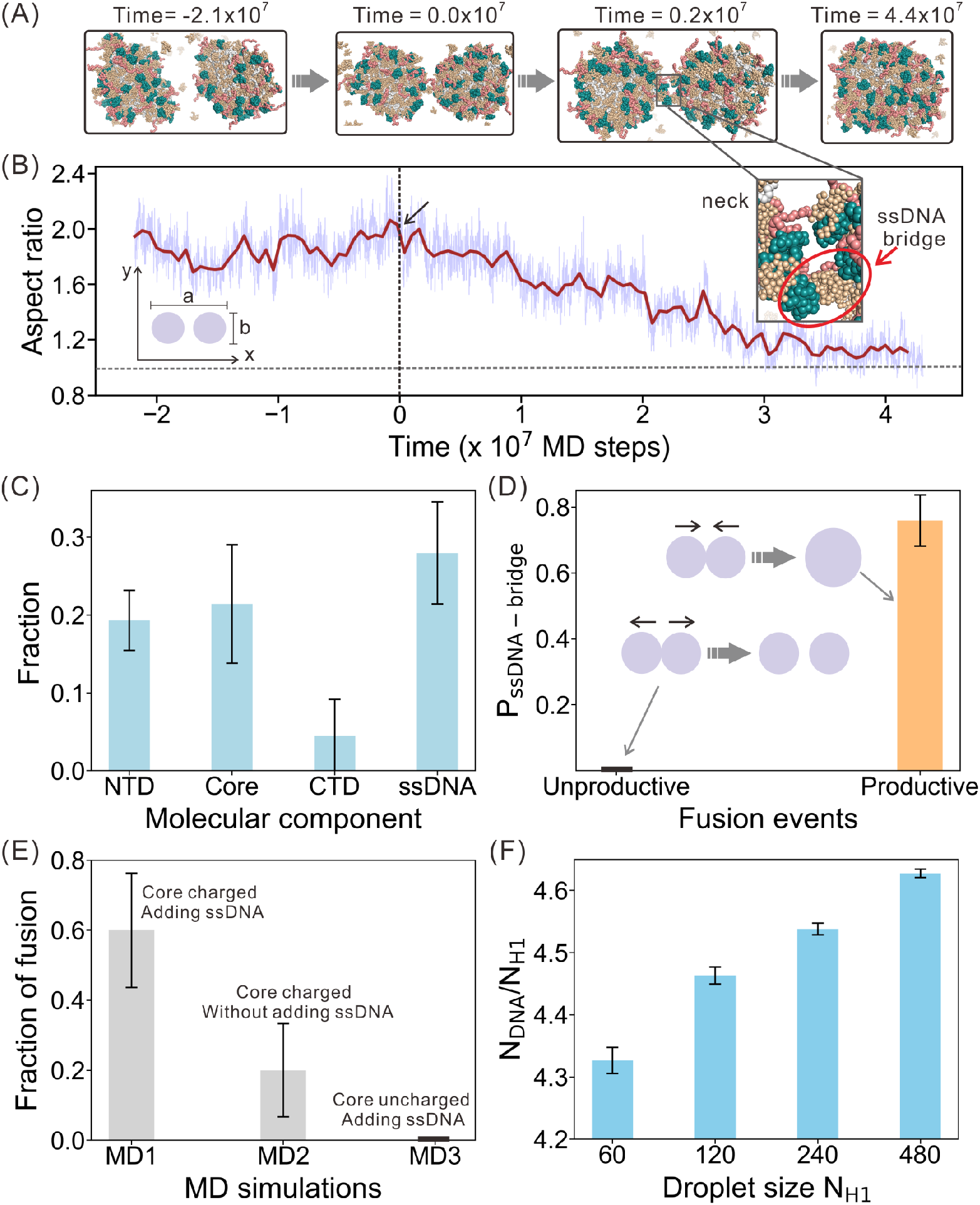
Molecular simulations of droplet fusion. (A) The snapshots along a representative trajectory of successful fusion event. The ssDNA-bridge in a neck-like region is highlighted. (B) The aspect ratio of the fusing droplets as a function of simulation time for the fusion event shown in panel (A). To establish a reference time point for the fusion process, the time step at which the two droplets initially came into contact was set as zero. The aspect ratio is given by the ratio of the longest axis (a) to the shortest axis (b) in the xy-plane. (C) Fraction of different components in the neck-like region formed by the fusing droplets. (D) Probabilities of ssDNA-bridge formation in the unproductive and productive fusion events. The probability in the unproductive fusion events is zero and labelled by a black bar. (E) Fraction of trajectories successfully captured the fusion events in three sets of simulations. The fraction in MD3 is zero. (F) Molar ratio of the ssDNA and protein in the droplets with different sizes. The droplet size is characterized by the number of H1 chains (N*_H_*_1_) contained in the droplet.

As depicted in Figure 3A, once the two droplets initially made contacts, a neck-like contacting zone emerged. The presence of this neck-like structure is crucial for understanding the molecular mechanism of droplet fusion. To examine the composition of this region, the fractions of different molecular components were analyzed, which showed that the neck region was rich in NTD, core domain and ssDNA (Figure 3C), whereas the CTD is rare. Interestingly, the simulation trajectory showed that the fusion events are intrinsically stochastic (Fig. S2). Even though two droplets come close and form contacting zone, it may either evolve to a fused droplet as shown in Fig. 3A (productive) or spontaneously separate again (unproductive). Remarkably, the productive fusion event relies on the formation of a ssDNA bridge wherein one ssDNA simultaneously forms contacts with two NTD/core domains from different fusing droplets. For the productive fusion events, the probability of observing the ssDNA-bridge in the contacting zone of the sampled snapshots is larger than 0.7, whereas the probability is zero for the unproductive fusion events. Based on these observations, it can be inferred that the ssDNA bridged interactions between the fusing droplets play a crucial role in facilitating subsequent coalescence.

To further investigate the key role of ssDNA-bridge in droplet fusion, we conducted two additional sets of simulations (Figure 3E). In the first set of simulations (MD2), there is no free ssDNA molecules being added to the simulation box. In comparison to the simulation results shown above (MD1), we observed a noticeable decrease in fusion ability. Out of the ten simulations conducted, only two successfully captured fusion events (Figure 3E, Figure S1). This decrease in fusion ability suggests again that the presence of free ssDNA molecules, which can accumulate onto the droplet surface, may play an important role in promoting efficient droplet fusion. The removal of these molecules appears to hinder the formation of ssDNA-bridge as discussed above. In addition, the presence of free ssDNA may also neutralize the surface charges of the droplets and therefore weaken the electrostatic repulsion during the approaching phase of the fusion event. In the second set of simulations (MD3), the charges of the residues in the core domain are neutralized. The structured core domain, which is the most prevalent component on droplet surface contains multiple positively charged residues and it is the main participant of the ssDNA-bridge. We observed that none of the trajectories captured droplet fusion event among the ten simulations (Figure 3E, Figure S1), which highlights again the crucial role of ssDNA-bridge formed in the contacting zone in promoting efficient droplet fusion and growth.

The above results clearly demonstrate that the ssDNA-bridge coupled kinetic control plays a crucial role in the droplet fusion and growth process. Furthermore, we observed that the preferred stoichiometry of the droplets depend on their sizes. For example, the molar ratio between ssDNA and H1 is *∼* 4.3 in the small droplet used in the above fusion simulations, which is much lower than the initial ratio (5.0). After the fusion of the two small droplets in the MD1 simulations, the molar ratio of the droplet becomes larger than 4.4 (Figure S3), which may suggest that larger droplet prefers a stoichiometry with higher ssDNA content. As a further characterization of the size dependence of the droplet stoichiometry, we performed additional droplet simulations, which generated the droplets with various sizes (See Methods section for more details). In line with the above observations, the results showed that as the droplets grew in size, the molar ratio of between ssDNA and H1 in the droplets tends to increase (Figure 3F). Such droplet size dependence of the stoichiometry suggests that the preferred molar ratio of the ssDNA and protein for droplet fusion changes with time, demonstrating again the key role of kinetic control in the droplet growth.

### Microstructure of condensates is size-dependent

The interactions between the biomolecules within the condensate dictate its microstructure and biological function.^70^ Therefore, we also performed a detailed characterization of the droplet microstructure formed by H1 and G4-ssDNA. For the droplet (N*_H_*_1_=40) used in the fusion simulations, we calculated the radial distribution function (RDF) of the mass density. The results revealed a core-shell-like structure of the droplet (Fig. S4). Specifically, the NTDs and core domains of H1 form a shell-like arrangement encasing a core composed of CTDs and ssDNA molecules, as also illustrated by the structural profile of the droplet.

To further investigate whether such core-shell-like structure is maintained for larger droplets, we also conducted structural analysis on the droplets of different sizes generated by the above droplet simulations. In the case of the droplet with N*_H_*_1_=60, a core-shell structure similar to the one observed above within a smaller droplet (N*_H_*_1_=40) was identified as shown by the droplet structural profile and spatial distribution of different H1 domains (Figure 4A-C, left). However, as the droplet increases in size, distinct structural changes become evident. The primary structural transformation observed during droplet growth involves an increased condensation of NTD and core domains towards the interior of the droplet (Figure 4A-C, middle). Additionally, as the droplets continue to enlarge, further microphase separation occurs, leading to additional structural variations. For example, in the case of a large droplet with N*_H_*_1_=480, the NTDs and core domains adopt a hollow-like microphase structure (Figure 4A-C, right), which is in contrast to the clustered substructure observed in a droplet with N*_H_*_1_=240. Furthermore, in a significantly larger droplet (N*_H_*_1_=960), these molecular components can undergo distinct microphase separation, resulting in the formation of various NTD/core clusters within the interior of the droplet (Figure S5). These findings suggest that the condensed structure within the droplets may exhibit a size-dependent nature. In other words, as the droplet size increases, the structural organization and behavior of the biomolecular components can undergo significant changes.

**Figure 4.**
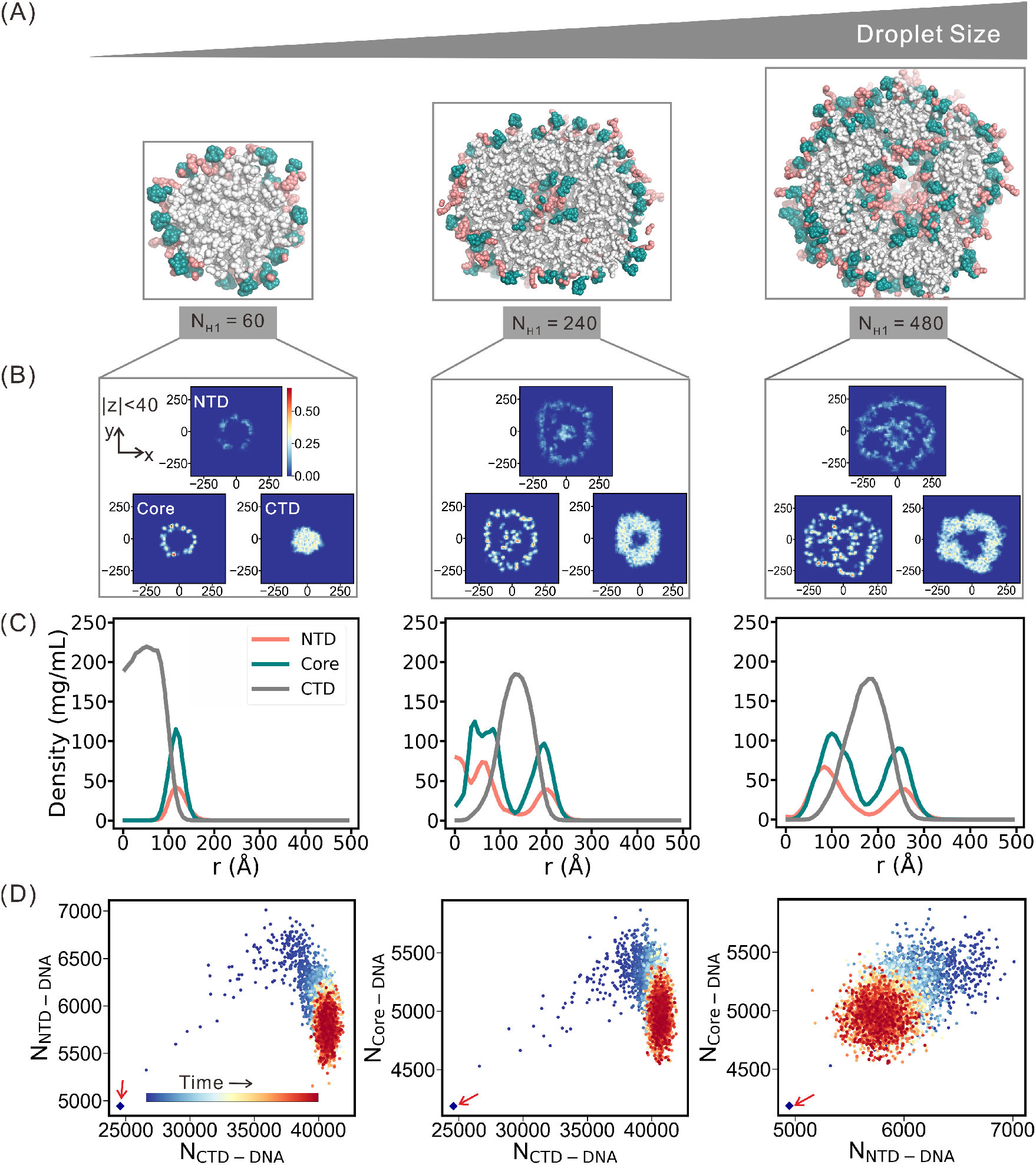
Size-dependence of droplet microstructure. (A) Structural profiles for three representative conformations of droplets with different sizes (N*_H_*_1_= 60, 240, and 480). Only the H1 molecules are displayed to ensure a clear visualization. (B) Distribution probabilities of the three domains of H1 within a slice of the droplets (*|*z*|≤*40). (C) Radial distribution of the mass density for the three domains of H1. (D) The trajectory of droplet simulations with the droplet size of N*_H_*_1_=480 projected along the reaction coordinates (N_CTD_*_−_*_DNA_, N_NTD_*_−_*_DNA_), (N_CTD_*_−_*_DNA_, N_NTD_*_−_*_DNA_), and (N_CTD_*_−_*_DNA_, N_NTD_*_−_*_DNA_). Here N_CTD_*_−_*_DNA_, N_NTD_*_−_*_DNA_, and N_core_*_−_*_DNA_ describe the number of formed contacts between ssDNA and the three domains of H1 within the droplet. The starting point is indicated by red arrow.

The above size dependence of the droplet microstructure can be understood based on the maximization of the electrostatic contacts between the highly charged CTDs and ssDNA. According to the sequence feature of H1, the CTD is highly charged, and therefore the contact formation of CTD with the oppositely charged ssDNA is energetically favored. For the droplet with small size, such a energetic requirement can be easily satisfied when the CTD and ssDNA are buried in the interior of the droplet, whereas the NTD and core domain are exposed to the droplet surface. However, with the growth of the droplet size, the CTD and ssDNA cannot occupy the full interior of the droplet. In this case, the NTD and core tend to form clusters in the interior of the droplet to maximize the favorable CTD-ssDNA contacts and therefore minimize the energetic frustration. ^47,71,72^ The increasing of the contact number between the CTD and ssDNA during the simulations of the droplets with different sizes (Figure 4D and Figure S6) clearly demonstrate the above energetic rule for the formation of droplet microstructure.

### Quadruplex folding promotes LLPS by increasing the multivalency and strength of protein-DNA interactions

The three-dimensional structures of protein and nucleic acid have the potential to affect the behavior of phase separation.^70^ In this section, we focused on investigating the influence of DNA structure on LLPS by comparing the phase separation behaviors induced by folded and unfolded G4 structures. For simplicity, we referred to the folded and unfolded G4-ssDNA as G4-F and G4-U, respectively, throughout this study. First we conducted slab simulations at 300K for the ssDNA-H1 mixture with the same molar ratio as the above simulations. For both the G4-F and G4-U systems, two distinct phases were observed, i.e., dilute phase and liquid-like condensed phase (Figure 5A,B), suggesting their capability of undergoing phase separations. However, compared to the G4-U system, the G4-F system demonstrated lower saturation concentrations (C*_sat_*) at the dilute phase but higher concentration at the condensed phase (Figure 5C). Such results imply that the system with the ssDNA folded into G4 structure is more prone to undergo phase separation, which is consistent with experimental observations.^61^

**Figure 5.**
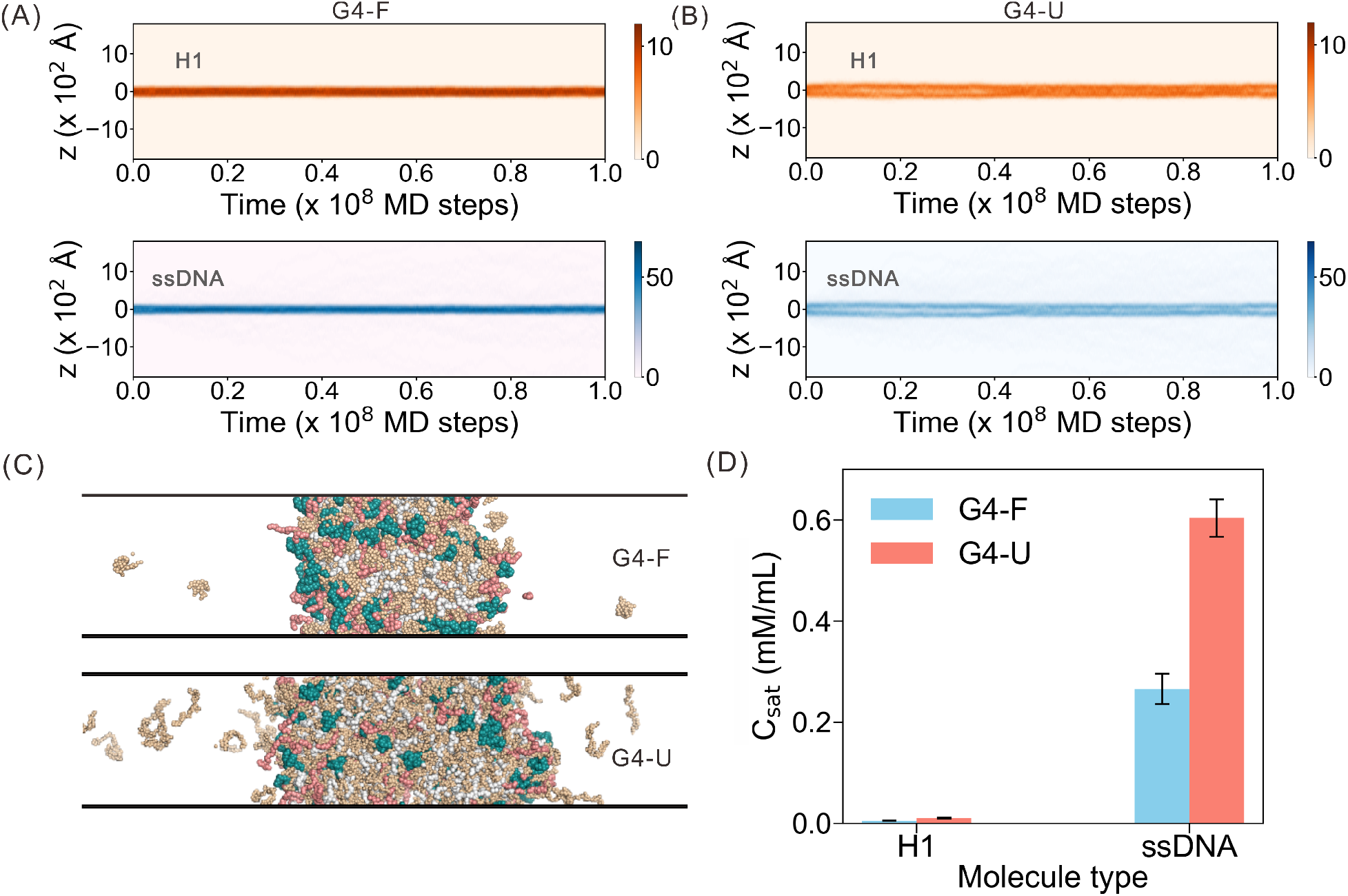
Effect of G4-ssDNA folding on phase separation. (A,B) Density profiles of H1 and ssDNA in the slab simulation trajectory for the systems G4-F (left) and G4-U (right) at 300K. (C) Representative snapshots of the slab simulations for the systems G4-F (upper) and G4-U (lower), respectively. (D) Saturation concentrations of the ssDNA and H1 estimated from the slab simulations for the systems G4-F (blue) and G4-U (red), which correspond to the concentrations in the dilute phase.

Next, to further understand the key role of G4 folding on the LLPS, we performed additional simulations for the G4-F and G4-U systems containing one H1 chain and 40 ssDNA chains, and analyzed the interaction pattern at the residue-level. We first calculated the contact probabilities between the residues of H1 and the nucleotides of ssDNA (Figure 6A). The contact map revealed that H1 in the G4-F system tends to interact with the G4 groove between two GGG-repeats, i.e., GGG_1_ and GGG_4_, which are positioned at the 5’-terminal and 3’-terminal respectively (Figure 6A and Figure S7). In contrast, no distinct binding pattern was observed for the G4-U system (Figure 6B). In addition, it was observed that the folded G4 exhibits a greater affinity for binding to both the NTD and core domain of H1, as illustrated by the increased contact probability (Figure 6C). These findings collectively indicate that the folding of G4 structure enhances the ability of ssDNA to bind to proteins. Furthermore, it was observed that a single chain of H1 can attach with more ssDNA molecules in the G4-F system compared to that in the G4-U system (Figure 6D). This observation suggests an increase in the interaction multivalency and strength between protein and ssDNA upon G4 folding. One possible reason for the enhanced interactions is that the folding of G4-ssDNA to a conpact structure increases the local charge density around the G4 grooves as contributed from the phosphate groups of the nucleotides, which therefore strengthens the ssDNA-H1 interactions. Particularly, the G4 groove between GGG_1_ and GGG_4_ is free of steric blocking by ssDNA loops, which can lead to the increased accessibility and contact probability of the positively charged domains of H1 as observed in Figure 6A.

**Figure 6.**
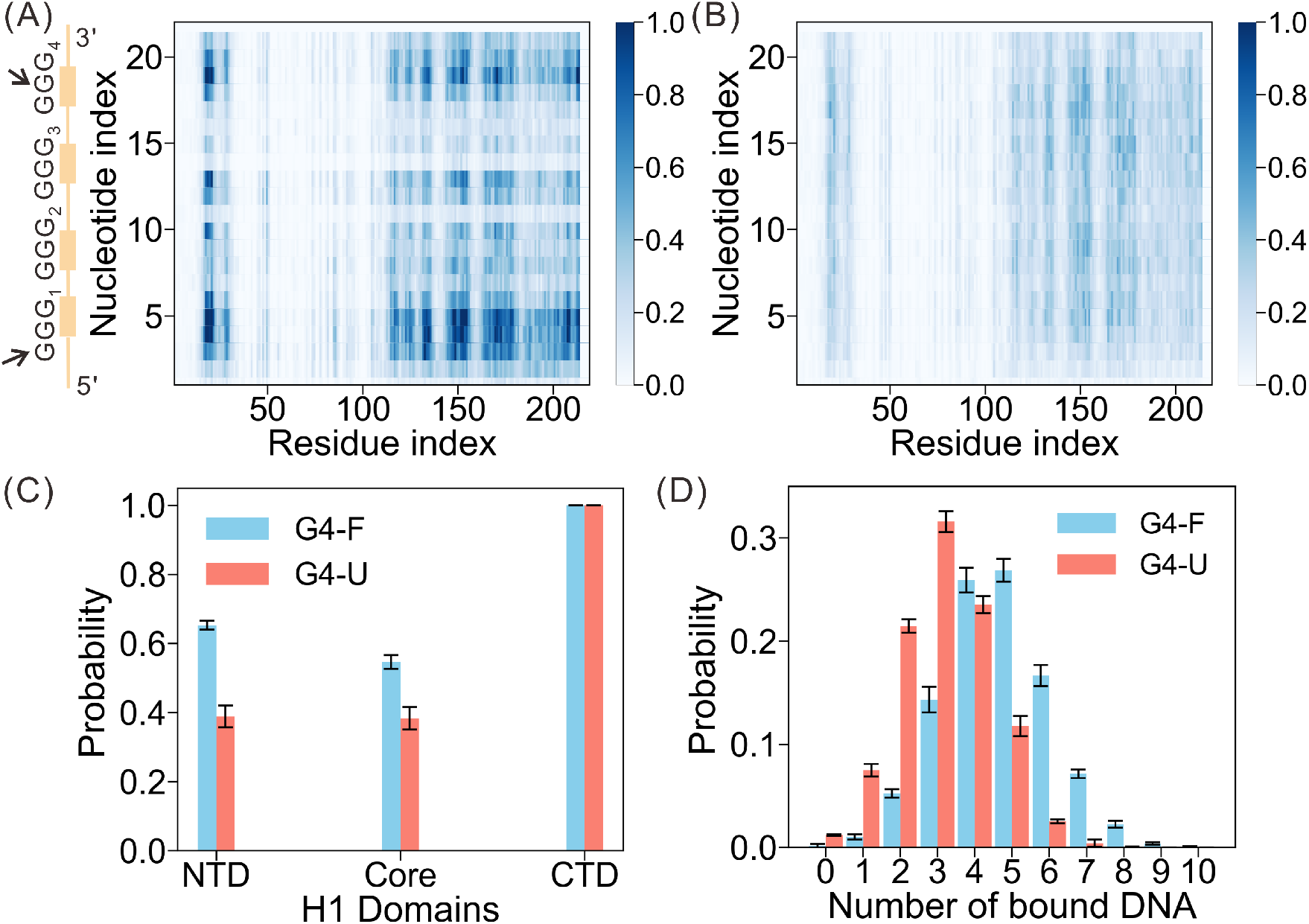
Effect of G4-ssDNA folding on the ssDNA-H1 interactions. (A,B) Probabilities of the contact formation between the residues H1 residues and ssDNA nucleotides for the systems G4-F (A) and G4-U (B). (C) Probabilities of ssDNA binding with the NTD, CTD, and Core domain of H1 for the systems G4-F (blue) and G4-U (red). (D) Distribution probability of the number of ssDNA molecules bound to one H1 chain for the systems G4-F (blue) and G4-U (red).

## Discussion

Understanding the mechanism of fusion dynamics is crucial in gaining insights into the growth of biological condensates and the related biological functions. It has been well established that the dynamics of droplet fusion are closely associated with the physical and material properties of the condensates, such as viscosity and surface tension, which in turn rely on the inter-molecular interaction strengths and the structural compactness of component macromolecules.^73^ In addition, recent studies revealed that the growth of biological condensates can also be regulated by modifying the droplet surface as a result of recruitment of additional biomolecules.^7,74–76^ For example, experimental investigation of P granule growth showed that the recruitment of MEG-3 protein clusters to the granule surface can impede its growth.^7^ In this work, we further revealed the critical role of the kinetic control in the droplet fusion by investigating the early stage of the ssDNA-H1 cocondensation. The molecular dynamics simulations directly demonstrated the stochastic collision and fusion processes of the droplets. The fusion event is productive only when an ssDNA simultaneously forms electrostatic contacts with the positively charged H1 core domains from the surfaces of two fusing droplets in the neck-like contacting zone. Diminishing such ssDNA-bridge by removing the free ssDNA or neutralizing the charge state of the core domain results in a significant attenuation of the fusion ability.

By adopting different shapes and arrangements, condensates can achieve specialized compartmentalization and provide distinct microenvironments for biochemical reactions to take place. It has been shown that these condensed structures are not always homogeneous spherical droplets.^2^ For instance, condensates composed of multiple components have been observed to adopt core-shell structures, as observed in paraspeckles, P granules, and stress granules.^77–79^ Additionally, certain condensates display hollow morphologies, such as the nuclear germ granules in Drosophila and nucleoprotein-RNA mixtures.^80,81^ Furthermore, under certain conditions, these condensate structures can undergo further microphase separation. An example is observed in the presence of type II topoisomerase enzymes, where wall-like substructures can emerge through microphase separation within euchromatic domains, which form via the phase separation of chromatin.^82^ Consistent with these observations, we identified an inhomogeneous structural feature within the condensate formed by H1 and ssDNA. Moreover, we have discovered that the microstructure of the condensate is dependent on its size. This indicates that the physical properties and organization of the condensate may vary based on its overall dimensions. In smaller condensates, the constituent molecules tend to form core-shell-like structures. However, as the condensate grows in size, the microstructure of the droplet core can undergo further phase separation. This microphase separation within the core region of the condensate highlights the dynamic nature of biology condensates and suggests a complex interplay between molecular interactions and the overall organization of the condensate.

Accumulated experimental reports have suggested that phase behavior of biomolecules can be altered by conformational changes. An example illustrating this observation is the nucleocapsid (NC) protein of HIV-1.^83^ In the absence of Zn^2+^, the NC protein is intrinsically disordered. However, upon the binding of Zn^2+^, it undergoes a conformational transition and folds into a structure that contains two zinc finger domains. Moreover, this conformational change was found to be crucial to initiate liquid-liquid phase separation (LLPS), leading to the formation of condensates. Another illustrative example in this context is the G4 structure studied here. The folding of G4 structures from ssDNA sequences has been found to play a pivotal role in promoting phase separation. Our simulations successfully reproduced this experimental observation.^61^ In addition, we observed that the folding of ssDNA into G4 structures significantly influenced the strength of interactions between the G4 structures and the H1 protein within the condensate. Furthermore, this change in interaction strength was shown to have a profound impact on the multivalency of specific interactions between proteins and DNA. Since the saturation of interaction multivalencies determines the final size of the condensate,^84^ the increased multivalency upon G4 folding allows the formation of larger condensates.

In summary, by performing molecular simulations, we studied the LLPS of H1 with a focus on the regulation by ssDNA that has the capability to fold into G4 structure. The phase separation process, including the dynamics of fusion events, was thoroughly characterized in our study (Figure 7). We showed that the fusion of droplets is a rather stochastic process and can be kinetically controlled. The formation of ssDNA-bridge between two fusing droplets represents the kinetic bottleneck step for the productive fusion event. Furthermore, our study revealed that the microstructure of the condensate is intricately linked to its size as a result of maximizing the electrostatic contacts. Microphase separation progressively occurs with the increasing of droplet sizes. In addition, the stoichiometry of the condenstate is also size dependent, and larger droplet prefers a stoichiometry with higher ssDNA content. Finally, our study suggests that the increase in multivalency and inter-molecule interaction strength between the H1 and ssDNA is a plausible mechanism responsible for the promotion of liquid-liquid phase separation (LLPS) by the folding of ssDNA into G4 structure,. These findings offer novel insights into the process of LLPS involving proteins and ssDNA, thereby enriching our understanding of biomolecular condensation.

**Figure 7.**
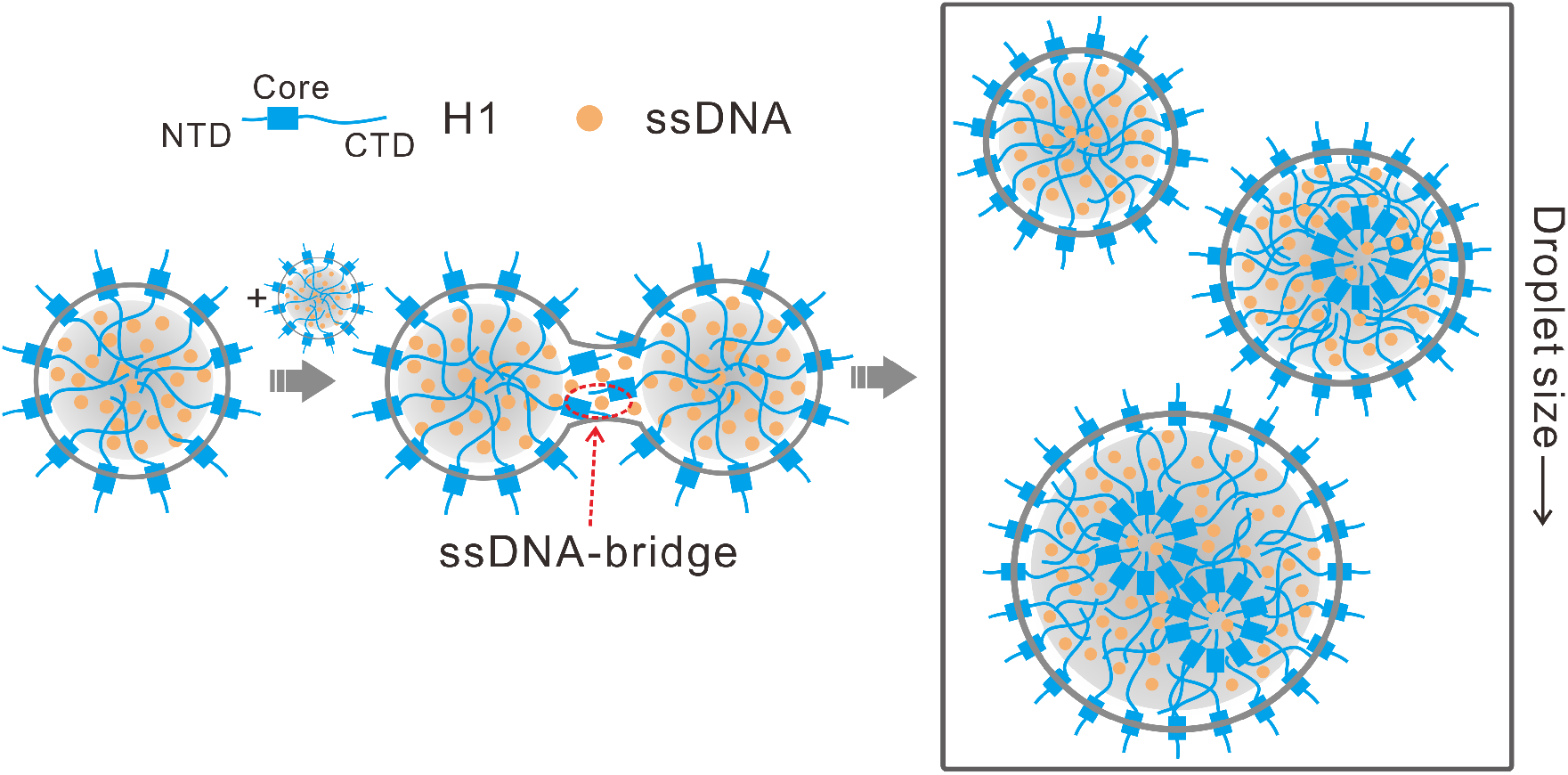
Schematic illustration showing the molecular mechanism of droplet fusion and growth. The ssDNA-bridge, which represents the bottleneck step for the productive fusion event, is highlighted by red circle.

## Materials and Methods

### Models

In this study, residue-resolved coarse-grained models were employed to characterize the protein condensation mediated by ssDNA. For protein, each residue was represented by one spherical particle centered at the C*α* position. The atomic interaction-based coarse-grained model with flexible local interactions (AICG2+)^72,85^ was used to describe the dynamics of folded domain of H1, while the hydrophobicity scale (HPS) model^86^ was employed to characterize the interactions involving the intrinsic disordered NTD and CTD. In addition, the three-site-per-nucleotide model (3SPN.2)^87^ was adopted for the ssDNA, in which each nucleotide was represented by three particles centered at the phosphate group, the sugar group and the base group, respectively. To describe the role of G4 structure in LLPS, an additional structure-based potential^88^ was added to restrain the ssDNA in the native structure. The Debye-Hükel type electrostatic potential was used to characterize the salt-concentration-dependent interactions of charged particles. The combination of AICG2+ and 3SPN models have been successfully used in the molecular simulations of the structure assembly, functional dynamics, and LLPS of the protein-DNA systems.^45,46,89^

### Simulation Details

The simulations were performed with the GENESIS package (version 1.7.1).^90–92^ The PDB structures with the entries 7K5Y and 1XAV were used as the references in building the structured potentials of the H1 and G4 fold, respectively.^93,94^ The mixing molar ratio of ssDNA and protein in all the simulations was 5.0 except otherwise stated, since the system was found to be most prone to undergo phase separation under this condition in experiments.^61^ The simulations were conducted in NVT ensemble except the ones with spherical boundary restraint. All the simulations were conducted using the Langevin dynamics with an inverse friction constant of 0.1 ps. The simulation temperature was 300K and the salt concentration was set to 150 mM. The time step of the MD simulations was 0.01 ps and the total simulation times of condensate dynamics and fusion dynamics were 1 *×* 10^8^ MD steps and 5 *×* 10^7^ MD steps, respectively.

To generate droplets of varying sizes, we utilized simulations employing a spherical potential to confine molecules within distinct spatial regions (droplet simulations), mimicking the formation of droplet-like structures. A specific number of H1 chains (N*_H_*_1_=60, 120, 240, 480 and 960) and the corresponding number of ssDNA chains (N*_DNA_*=300, 600, 1200, 2400 and 4800) were randomly placed in a spherical box. Initially, an equilibrium simulation of 1 *×* 10^7^ MD steps was performed. Subsequently, a productive simulation of 2 *×* 10^7^ MD steps was conducted, involving the enlargement of the spherical box radius. Such droplet simulations allowed for the examination of droplet formation and behavior under varied conditions.

In slab simulations, the system contained 60 chains of H1 and 300 chains of ssDNA, which is sufficient to demonstrate the different phase behaviors of H1 under the regulation of G4 folding. Each system was placed in a periodic box with a long z-axis (200*Å* × 200*Å* × 3600*Å*). The initial conformation was a condensed structure obtained from a “shrinking” simulation that squashed the box to 200*Å* × 200*Å* × 200*Å* in 1 *×* 10^6^ MD steps at 300K. Both slab simulations for G4-F and G4-U systems started from this conformation. In the case of G4-F, a structure-based potential was added to restrain the ssDNA in parallel G4 topology.^95^ Each slab simulations lasted for 1 *×* 10^8^ MD steps and the snapshots sampled during the last 5 *×* 10^7^ MD steps in each trajectory were used in the analysis.

During the clustering of H1 chains in the simulations of condensate formation, we estimated the contacts formed by proteins and ssDNA molecules for each snapshot. The H1 monomers were grouped into the same cluster when they interacted with the same ssDNA. A contact was considered to have formed when two beads were within a distance of 5 *Å*.

In the droplet fusion simulation, the neck structure was defined as the substructure formed by the two droplets from the moment of their initial contact to the point at which stable interactions were established. It was observed that productive fusion occurred when the distance between the two droplets, specifically the center of mass of the H1 chains in the droplets, was less than 240 *Å*. As a result, we determined that fusion was considered successful when the two droplets remained within a distance of 240 *Å* and did not dissociate for a specified duration of MD steps, which was set to 5 *×* 10^5^ in this study.

To examine the impact of the ssDNA-bridged substructure formed in the neck-like structure during fusion, we introduced a parameter H. This parameter was defined to characterize the interaction pattern of the ssDNA with the two droplets, utilizing a Hill function. Specifically, H was calculated as the average of *H*_1_ and *H*_2_, where *H_i_* (*i*=1,2) was given by *H_i_*= 1*/*(1 + *e^−(Qi−Qc)/σ)^*. In this equation, *Q_i_* represents the contact number between ss-DNA and the droplet *i*, which quantifies the extent of contact formation between the ssDNA molecules and the core domains within the neck region. *Q_c_* serves as a cutoff threshold for Q, dictating the formation of stable interactions. Meanwhile, *σ* is a parameter that modulates the shape of the Hill function. In this work, *Q_c_* and *σ* were selected as 5.0 and 1.0, respectively. With these defined settings, the resulting H values can effectively quantify the number of droplets to which a ssDNA molecule binds. H≈0.0 indicates that the ssDNA molecule did not bind to any droplets, whereas H≈0.5 suggests that the ssDNA molecule is binding to one droplet. On the other hand, an H value of *∼*1.0 indicates that the ssDNA molecule is binding to two droplets simultaneously, forming a ssDNA-bridge.

## Acknowledgement

The authors thank Zhi Qi and Cheng Li for insightful comments. This work was supported by National Natural Science Foundation of China [Grant no. 11974173], the grant of Wenzhou Institute, University of Chinese Academy of Sciences (WIUCASQD2021010 and WIUCASQD2022036) and Natural Science Foundation of Shandong Province (Grant no. ZR202102210546). The authors also thank the support of High Performance Computing Center of Nanjing University.

